# The GL Service: Web Service to Exchange GL String Encoded HLA & KIR Genotypes With Complete and Accurate Allele and Genotype Ambiguity

**DOI:** 10.1101/020099

**Authors:** Robert P. Milius, Michael Heuer, Mike George, Jane Pollack, Jill A. Hollenbach, Steven J. Mack, Martin Maiers

**Author notes:** **Communicating Author:** Robert P. Milius 3001 Broadway St NE, Suite 100 National Marrow Donor Program Minneapolis, MN 55413-1753 USA. **Conflicts of Interest:** There are no conflicts of interest.

## Abstract

Genotype List (GL) Strings use a set of hierarchical character delimiters to represent allele and genotype ambiguity in HLA and KIR genotypes in a complete and accurate fashion. A RESTful web service called Genotype List Service was created to allow users to register a GL String and receive a unique identifier for that string in the form of a URI. By exchanging URIs and dereferencing them through the GL Service, users can easily transmit HLA genotypes in a variety of useful formats. The GL Service was developed to be secure, scalable, and persistent. An instance of the GL Service is configured with a nomenclature and can be run in strict or non-strict modes. Strict mode requires alleles used in the GL String to be present in the allele database using the fully qualified nomenclature. Non-strict mode allows any GL String to be registered as long as it is syntactically correct. The GL Service source code is free and open source software, distributed under the GNU Lesser General Public License (LGPL) version 3 or later.

**Abbreviations:**

API: Application Program Interface
AWS: Amazon Web Services
CDISC: Clinical Data Interchange Standards Consortium
DaSH: Data Standard Hackathon
EMBL: European Molecular Biology Laboratory
EMDIS: European Marrow Donor Information System
ENA: European Nucleotide Archive
FDA: Food and Drug Administration
FHIR: Fast Healthcare Interoperability Resources
GL: Genotype List
GNU: GNU’s Not Unix
HL7: Health Level Seven, International
HLA: Human Leucocyte Antigen
HML: Histoimmunogenetics Markup Language
HTML: Hypertext Markup Language
HTTP: Hypertext Transfer Protocol
IMGT: ImMunoGeneTics
ISO: International Organization for Standardization
JDBC: Java Database Connectivity
JSON: Javascript Object Notation
KIR: Killer-cell Immunoglobulin-like Receptor
LGPL: Lesser General Public License
LSDAM: Life Sciences Domain Analysis Model
MHC: Major Histocompatibility Complex
MIRING: Minimum Information for Reporting Immunogenomic NGS Genotyping
MUG: Multilocus Unphased Genotype
N3: Notation 3
NCI: National Cancer Institute
NGS: Next Generation Sequencing
NMDP: National Marrow Donor Program
OID: Object Identifier
OWL: Web Ontology Language
PNG: Portable Network Graphics
QR Code: Quick Response Code
RAM: Random Access Memory
RDBMS: Relational Database Management System
RDF: Resource Description Language
REST: Representational State Transfer
SBT: Sequence Based Typing
SDK: Software Development Kit
SQL: Structured Query Language
SSO: Sequence Specific Oligonucleotide
SSP: Sequence Specific Primer
URI: Uniform Resource Identifier
URL: Uniform Resource Locator
XML: Extensible Markup Language

**Abbreviations:**

GL Service: A web service to exchange GL Strings

## 1. Introduction

Ambiguity in recording and reporting of HLA genotyping results continues to impact clinical decision support for donor/patient matching as well as understanding post-transplant outcomes research. Recommendations addressing this have been proposed [1]. These guidelines include listing all allele and genotype ambiguities at the highest resolution detected, documenting the typing method, and identifying the pertinent reference allele sequence database version used to perform the typing. To address the first part of these guidelines, a string format capable of fully representing HLA allele and genotype ambiguity was developed. This format, named Genotype List String (GL String), was developed by extending a proposed standard for reporting KIR genotyping data for use with HLA [2]. The GL String is parsed based on character delimiters that organize the alleles in terms of loci, alleles, lists of possible alleles, phased lists of alleles, genotypes, lists of possible genotypes, and multilocus unphased genotypes (MUGs).

An important goal of the GL String format is to separate the encoding of genotype data from the management and presentation of those data. GL Strings are potentially quite long and difficult to read, and are expected to proliferate rapidly in number. However, they are easily machine generated and parsed, and the work of creating and displaying them should be left to machines. The remaining challenge is one of exchanging the strings easily.

Toward this end, we have developed the GL Service, a web service that allows users to register GL Strings they have generated, and that returns a unique Uniform Resource Identifier (URI) [3] for each registered GL String. When provided with a URI, the GL Service will return a GL in a variety of formats. In this way, use of the GL Service improves the portability of genotyping results; URIs can be shared between parties, and identical GL Strings can be obtained from the GL Service. As described in greater detail in sections 2 and 3, the use of the GL Service as a hub for sharing genotyping results allows the identification of the pertinent reference allele database version for a given GL String, and establishes the equivalence of GL Strings across difference reference allele database versions.

The GL Service is currently available at https://gl.nmdp.org. A GL Service Explorer page for IMGT/HLA Database release version 3.18.0 is available at https://gl.nmdp.org/imgt-hla/3.18.0/explorer/. While an individual can use the Explorer to register GL Strings and receive URIs, and to dereference URIs and receive GLs, the GL Service is primarily intended to interact with other services and programs in an automated fashion. The GL Service code is open-source and available, and anyone may run a public or private version of their own GL Service. Here, we describe the current implementation of the GL Service, and the available libraries, tools and approaches for interacting with the service.

## 2. Methods

### 2.1. Genotype List Service

The Genotype List Service is implemented in Java in a modular fashion. It was designed to be secure, scalable, and persistent. Web services based on REpresentational State Transfer (REST) architectural style [4,5] with HTTP content negotiation were developed employing a Java library that manages genotyping results. Resources are identified with a structured URI.

The service makes reference to the IMGT/HLA Database [6], and allows aggregation from alleles to lists of alleles, haplotypes, genotypes, lists of genotypes and multi-locus unphased genotypes. RESTful web service APIs are provided for programmatic storage and retrieval from other software. The service supports content negotiation, and can be configured with a nomenclature and allele reference database version in either strict or non-strict modes.

Public services include creating and retrieving these objects. While REST allows four CRUD operations (create, retrieve, update and delete), the GL Service only provides create (POST) and retrieve (GET) operations. No updating or deleting of registered resources is permitted.

### 2.2. Implementation

The core model of the Genotype List Service follows directly from the Genotype List String (GL String) grammar described previously [2]. An abstract superclass Genotype List resource provides URI and GL String representation data properties as defined in Table 1.

**Table 1.**
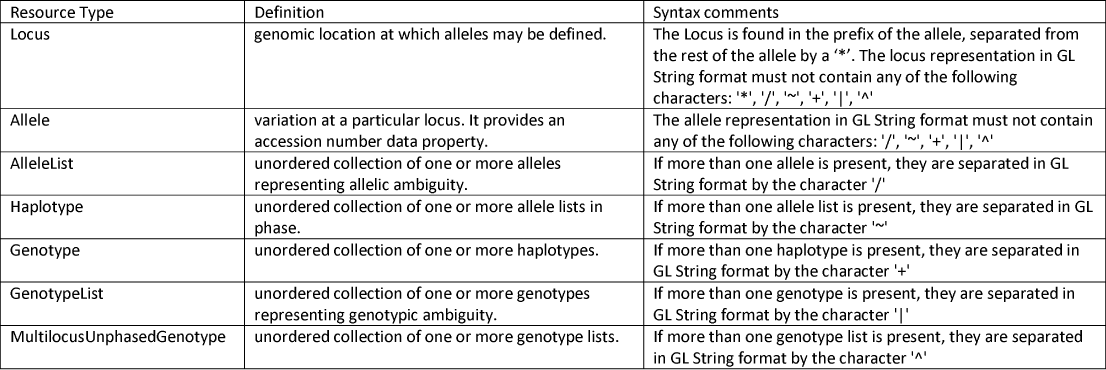

The core model is implemented as a Java library, defined as an ontology in OWL/RDF format [7,8], and described in XML Schema [9,10], in both object graph form and linked data form using XLink [11].

The Genotype List service builds on this core model and provides Java APIs for registering GL resources given a GL String and for retrieving GL resources given a URI identifier. An instance of the service is composed of various implementation modules using dependency injection. This allows the same code base to be deployed in various configurations according to operational environment characteristics and/or performance considerations.

The Genotype List service can be configured to use several persistence back ends. A relational database persistence implementation module supports both MySQL [12] and Oracle [13]. Additional No-SQL implementation modules offer potential read performance and horizontal scalability beyond that of a RDBMS. Finally, a RAM-only cache persistence implementation module is provided for functional testing, which runs as part of a continuous integration build process.

### 2.3. Availability

The GL Service is available as free open source software, with the source code distributed under the GNU Lesser General Public License (LGPL) version 3 or later.

Multiple versions of the GL Service, the API Explorer, and the liftover service configured with the IMGT/HLA Database reference alleles in strict mode are available at https://gl.nmdp.org.

The GL resource ontology in OWL/RDF format and in XML Schema, both in object graph form and in linked data form, are also available.

Source code and documentation can be found on https://github.com/nmdp-bioinformatics/genotype-list.

## 3. Results

### 3.1. Genotype List Service

#### 3.1.1. RESTful web service APIs

The GL Service advertises its functionality via web service APIs in the REST architectural style [4,5]. An endpoint URL is provided for each GL resource type. HTTP requests to these endpoints perform operations based on HTTP verbs (i.e., HTTP GET and HTTP POST).

To retrieve information about a GL resource identified by a URI, a client uses HTTP GET (e.g. HTTP GET https://gl.nmdp.org/imgt-hla/3.18.0/allele-list/0). If the service does not know anything about a particular URI, an HTTP 404 Not Found error is returned.

To register a new GL resource described by a string in GL String representation, a client uses HTTP POST to the appropriate endpoint (e.g. for allele lists, HTTP POST https://gl.nmdp.org/imgt-hla/3.18.0/allele-list) with the GL String representation in the body of the request. On success, the service will return a response with a HTTP 201 Created status code, the new URI in the Location header field, and the GL String representation of the newly created GL resource in the response body (as text/plain media type). On failure, an HTTP 400 Bad Request error is returned.

HTTP requests using other HTTP verbs (PUT, PATCH, DELETE) are not supported by the RESTful web service APIs.

#### 3.1.2. Content negotiation

For convenience, each GL resource is available in several different representations (GL String, HTML, XML, JSON, OWL/RDF, N3 and QR Code). The GL Service uses content negotiation to return the representation that best meets the needs of the client (or user agent). The user agent expresses its preferences by providing an Accept HTTP header that lists acceptable media types and associated quality factors.

The GL service will return text/plain media type by default for command line tools such as curl or wget, text/html media type for web browsers such as Firefox or Chrome, and RDF/XML media type for semantic web browsers and reasoners (e.g. Protégé [14] or Jena [15]).

In the case that content negotiation may not work as intended, the GL service also supports the use of file extensions in the query URL (e.g., .json, .rdf, .n3, .xml, .xml-xlink). For example, to force text/n3 media type in a web browser user agent, use the .n3 file extension in the query URL. (e.g., HTTP GET https://gl.nmdp.org/imgt-hla/3.18.0/allele-list/0.n3).

The GL service also provides the generation of a QR Code PNG image encoded with the URI identifier of the requested resource by use of the .png file extension. This could provide for example a small 2D barcode representing the full allelic and genotypic ambiguity of a typing result across several loci (i.e., a multilocus unphased genotype) in a small space on a typing report. For example https://gl.nmdp.org/imgt-hla/3.18.0/allele-list/0.png will return an image of a QR Code that redirects the scanning tool to the URI representing HLA-A*01:01:01:01/HLA-A*01:01:01:02N.

See Supplementary Material for more examples of different formatted views of the same genotype.

#### 3.1.3. IMGT/HLA Database release versions

To support our primary use case, where the IMGT/HLA Database [6] provides loci and alleles used to compose higher-level GL String representations, an instance of the GL Service can be configured with a specific release version of an allele reference database.

All of the loci and alleles provided by this database are registered as the service initializes. For the most recent version of the IMGT/HLA Database (Release 3.20.0, 2015-04-17), this consists of 35 loci and 13023 alleles.

#### 3.1.4. Strict/non-strict mode

Complementary to the nomenclature configuration parameter, the GL Service can be configured in either strict or non-strict mode. Strict mode requires alleles used in the GL String to be present in a particular version of a reference allele database (e.g., IMGT/HLA version 3.20.0). Non-strict mode allows any GL String to be registered as long as it is syntactically correct.

In the default non-strict mode, the GL Service will create new loci and alleles as necessary to register higher-level GL resources. For example, if an allele list represented in GL String format by “HLA-A*01:01:01:01/HLA-Z*99:99:99:99” was registered against a non-strict-mode instance of the GL Service, a new HLA-Z locus and HLA-Z*99:99:99:99 allele would be created in order to correctly represent the allele list.

In strict mode instances of the GL Service, pre-populated with specific reference allele lists, this default behavior is overridden, so that GL resources that do not adhere to the allele database release version in question cannot be registered. Any attempts to register an invalid GL resource will result in an error.

In strict mode, the namespace includes the name (e.g., “imgt-hla”, “kir”) and version number of the reference database. In non-strict mode, the name and version of the reference database are not included in namespace of the service. For example, the service found at https://gl.nmdp.org/imgt-hla/3.18.0 only accepts alleles found in version 3.18.0 of the IMGT/HLA Database. Using strict mode, a new instance of the service must be created for each reference database release. In strict mode, nomenclatures may not be mixed (e.g., HLA and KIR) within a GL String unless the nomenclature databases have been preloaded into the instance.

In strict mode, each allele must be represented as a complete object, including the full locus name with prefix. For example, HLA-A*01:01:01:01 is acceptable, but A*01:01:01:01 is not. The latter would be acceptable in non-strict mode, but this mode loses the ability to validate the allele. In this case, validation would be a responsibility of the creator of the GL-String and/or the consumer of the GL String referenced by the URI.

In strict mode, IMGT/HLA G groups are recognized as accepted HLA nomenclature (e.g., HLA-A*01:01:01G), but P groups are not. G groups are registered as alleles, although in reality they are in fact lists of alleles. They are accepted in strict mode as a courtesy to those who wish to exchange validated GL Strings containing G groups. In the future, a separate service may be developed to translate GL Strings containing G groups into IMGT/HLA Database version specific lists of alleles.

At the time of this writing, 63 strict GL Services are available representing IMGT/HLA Database versions from 1.5.0 to 3.18.0. Development of services for 3.19.0 and 3.20.0 are in progress.

### 3.2. Client libraries and tools

Since RESTful web service APIs are based on HTTP, the service endpoints are accessible from web browsers, browser-based RESTful web service clients (e.g. RESTClient [16], REST Easy [17]), and from command-line HTTP clients.

For example, to register a GL String via HTTP POST, the `curl` command may be used:

~~~
$ curl -v --header “content-type: text/plain” \
    --data “HLA-A*01:01:01:01/HLA-A*01:01:02+HLA-A*02:01:01:01” \
    -X POST https://gl.nmdp.org/imgt-hla/3.18.0/genotype
~~~

This above command will register the GL String and return the URI in the Location field of the response. In this case the returned URI is https://gl.nmdp.org/imgt-hla/3.18.0/genotype/u

To retrieve the contents of this URI, the wget command may be used. For example,

~~~
$ wget -q -O - https://gl.nmdp.org/imgt-hla/3.18.0/genotype/u
~~~

returns the GL String HLA-A*01:01:01:01/HLA-A*01:01:02+HLA-A*02:01:01:01

The GL Service project provides functionality beyond these generic REST clients, such as an interactive API Explorer, client libraries, and command line tools.

#### 3.2.1. API Explorer

An interactive web tool demonstrating the GL Service API is available at https://gl.nmdp.org/imgt-hla/3.18.0/explorer/.

This tool is not meant to be for production use, but rather provides an educational interface to the service, showing how GL Strings can be registered in the service, and how URIs can be dereferenced to retrieve GL Strings. The Explorer can also be used to show how different content types can be retrieved and display their output.

Figure 1 shows the interface to the API Explorer. The user selects the method that describes the object that is being registered (Figure 2). In this example, the GL String represents a genotype consisting of a list of two alleles and a single allele: HLA-A*01:01:01:01/HLA-A*01:01:02+HLA-A*02:01:01:01. The string is entered into the text entry box on the right and the HTTP POST button is clicked. The response from the service is displayed at the bottom of the page showing the Location of the resource in the form of a URI. In this example the URI is https://gl.nmdp.org/imgt-hla/3.18.0/genotype/u (Figure 3).

**Figure 1.**
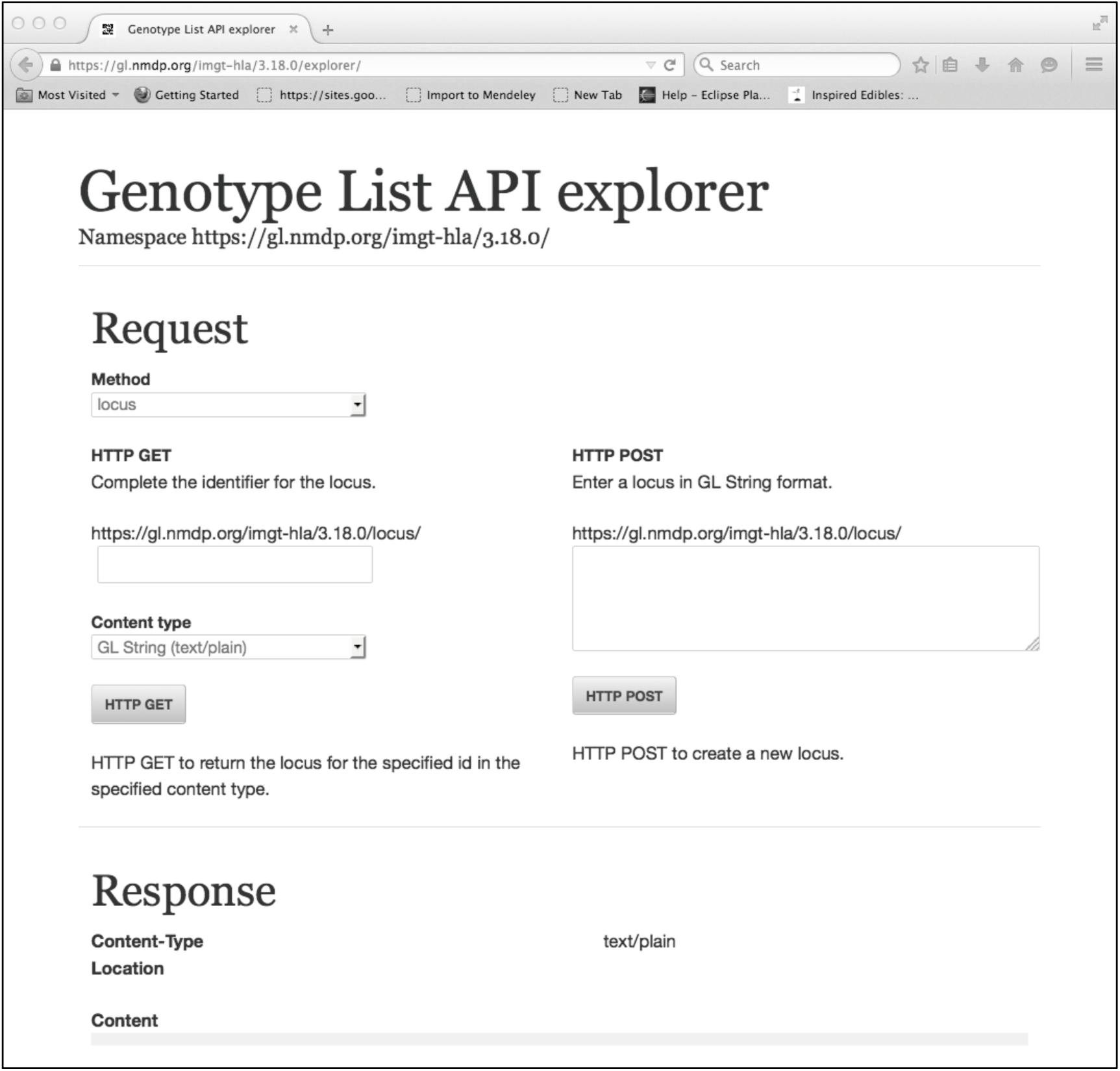
Interface of API Explorer found at https://gl.nmdp.org/imgt-hla/3.18.0/explorer/

**Figure 2.**
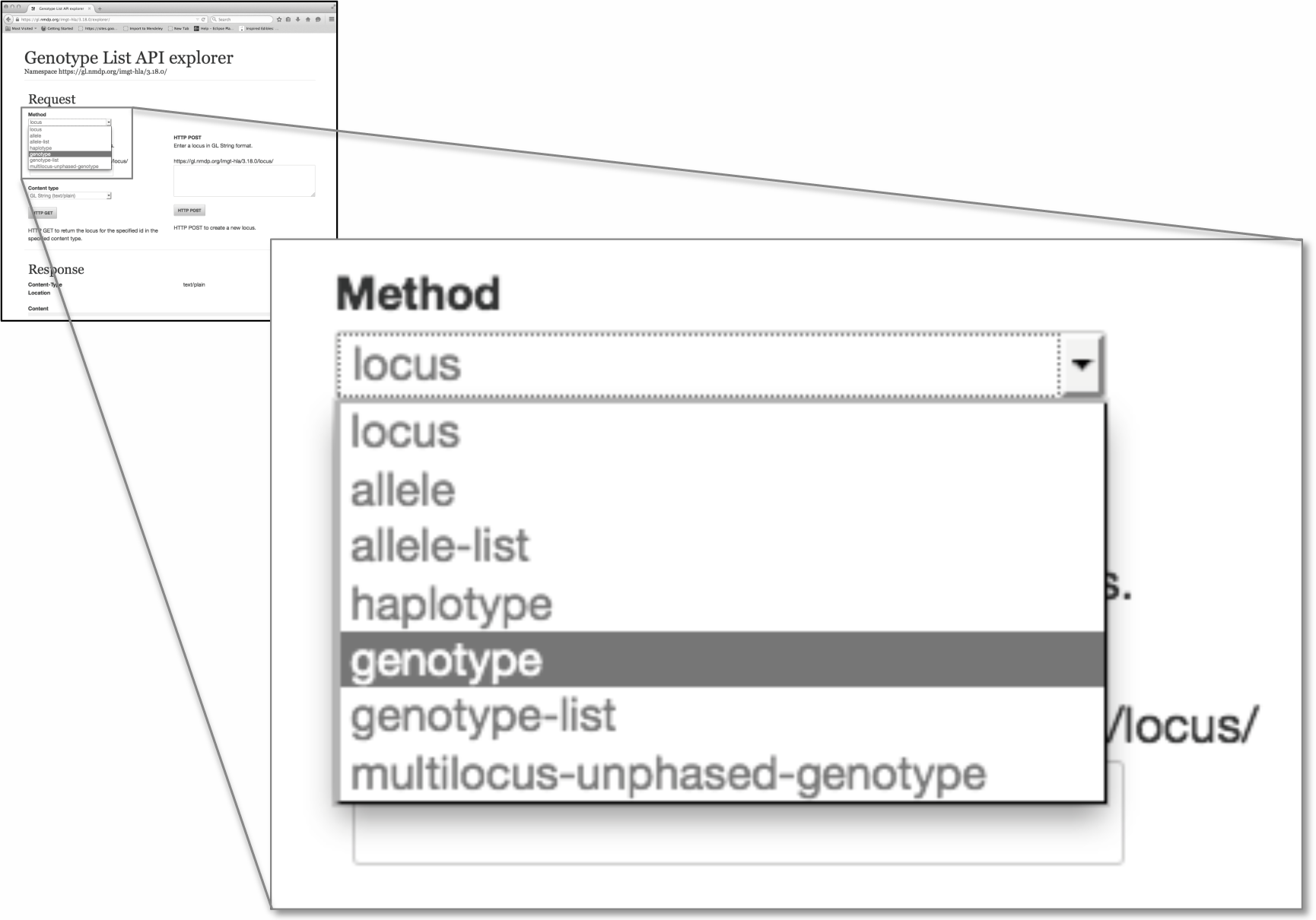
Selecting method in the API Explorer. The main panel is an exploded version of the Method selection box for the smaller panel, which is the entire view of the API Explorer. Here the Method selected is genotype.

**Figure 3.**
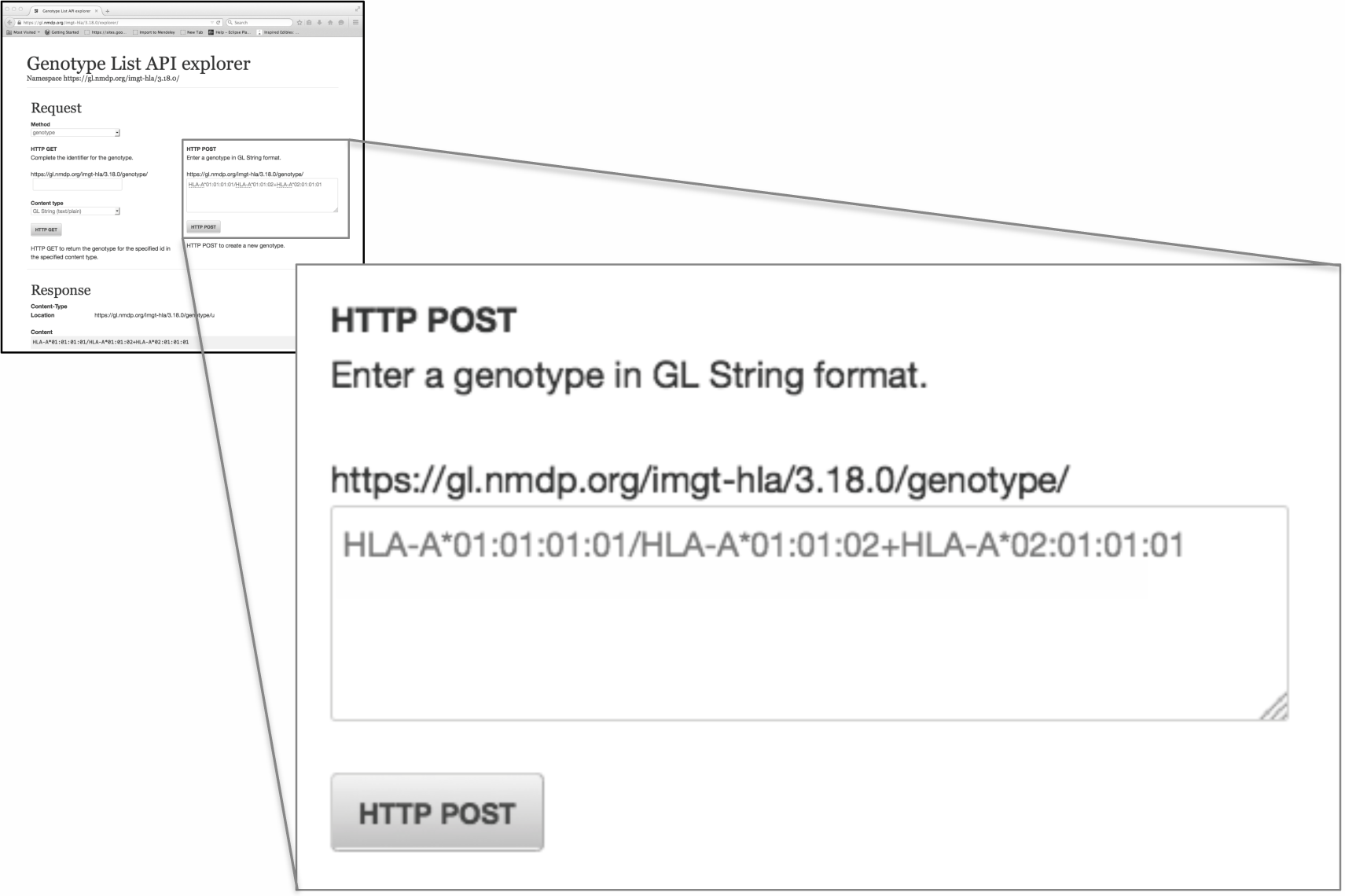
Using HTTP POST to register a GL String. The main panel is an exploded version of the HTTP POST entry box for the smaller panel, which is the entire view of the API Explorer. Here the GL String representing the genotype HLA-A*01:01:01:01/HLA-A*01:01:02+HLA-A*02:01:01:01 is entered.

The URI can then be used to retrieve the genotype in GL String format, or any one of other formats including HTML, XML, JSON, OWL/RDF, N3 and QR Code (Figure 4). Clicking the HTTP GET button retrieves the object and displays it in the selected content type. Figure 5 shows the response of the service in GL String format. Figure 6 shows the result of a request for the object in QR Code format. Scanning this QR Code with a scanning application found on smart phones will result in the URI opening in a web browser, showing the object in HTML format, which can be drilled deeper into the structure by following links (Figure 7).

**Figure 4.**
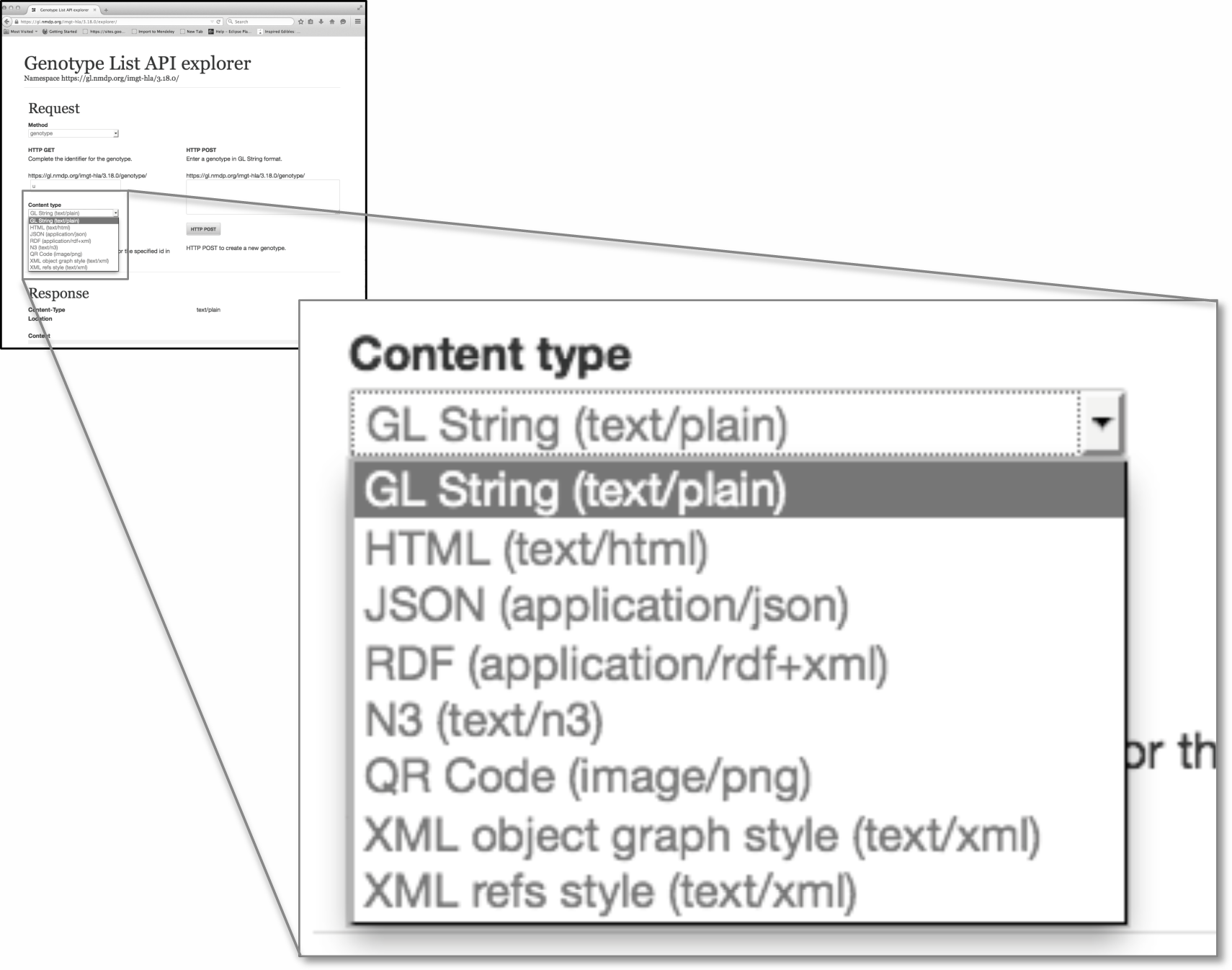
Selecting content type in the API Explorer. The main panel is an exploded version of the Content Type selection box for the smaller panel, which is reflects the entire view of the API Explorer. This is used to select the format when retrieving the dereference URI. In this case, GL String is selected.

**Figure 5.**
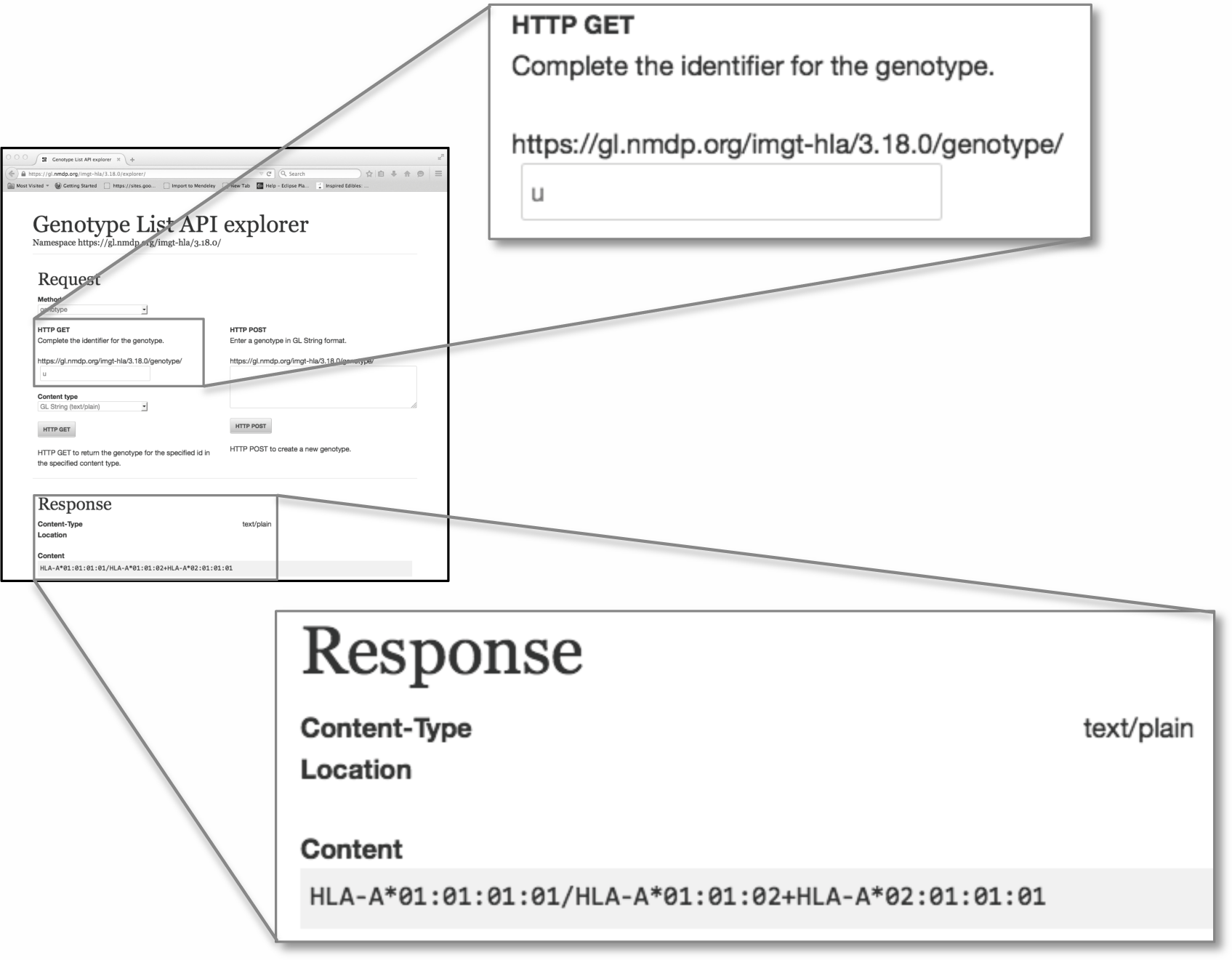
Using HTTP GET to retrieve a GL String. The URI for the genotype is completed using the data entry box. The Response is the GL String of the genotype returned after the GET command is sent.

**Figure 6.**
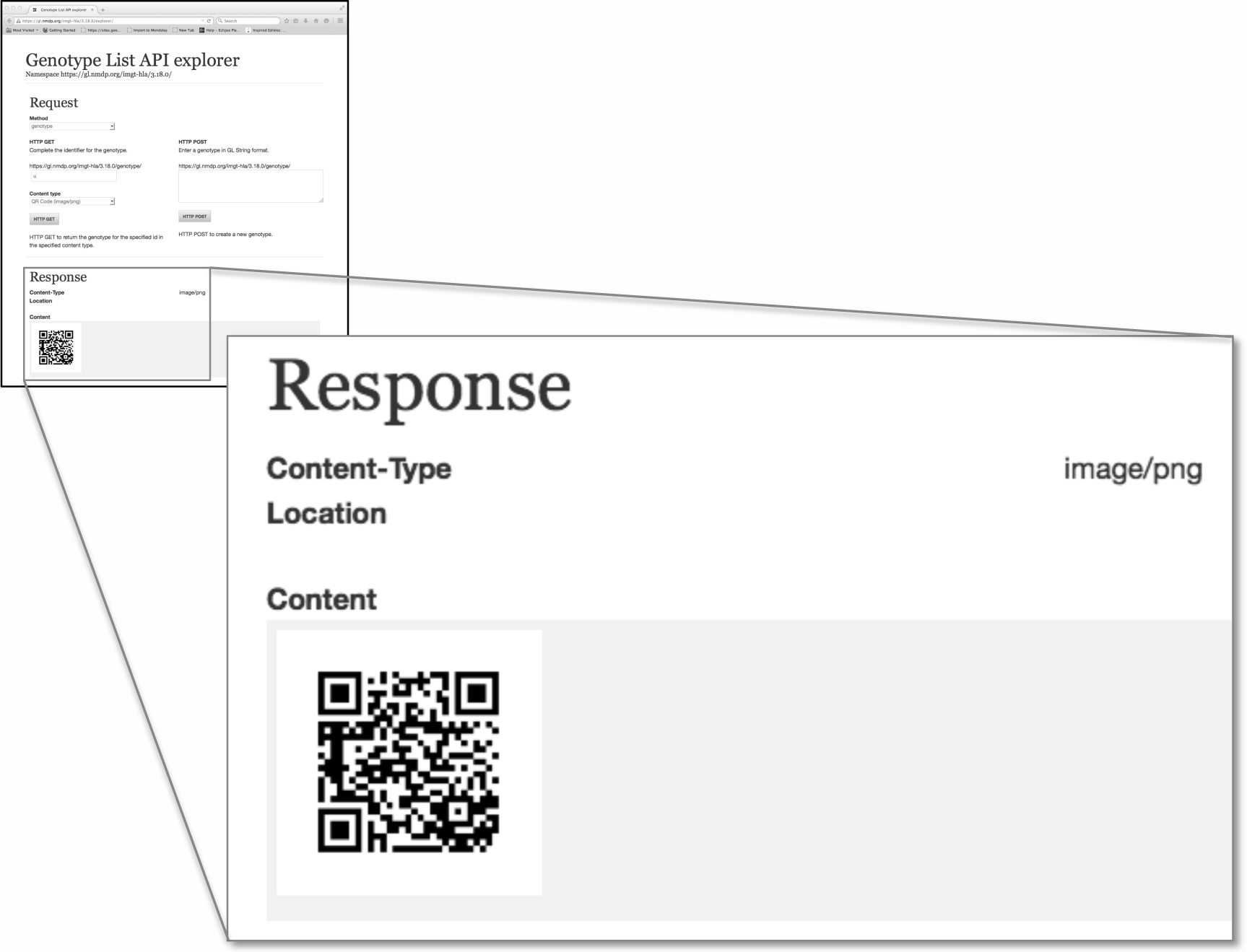
Using HTTP to GET a QR Code. The Response when QR Code is selected from Content Type. When scanned, the URI representing the GL String is returned. If the scanner is configured to open a web page, it will return the HTML representation of the URI.

**Figure 7.**
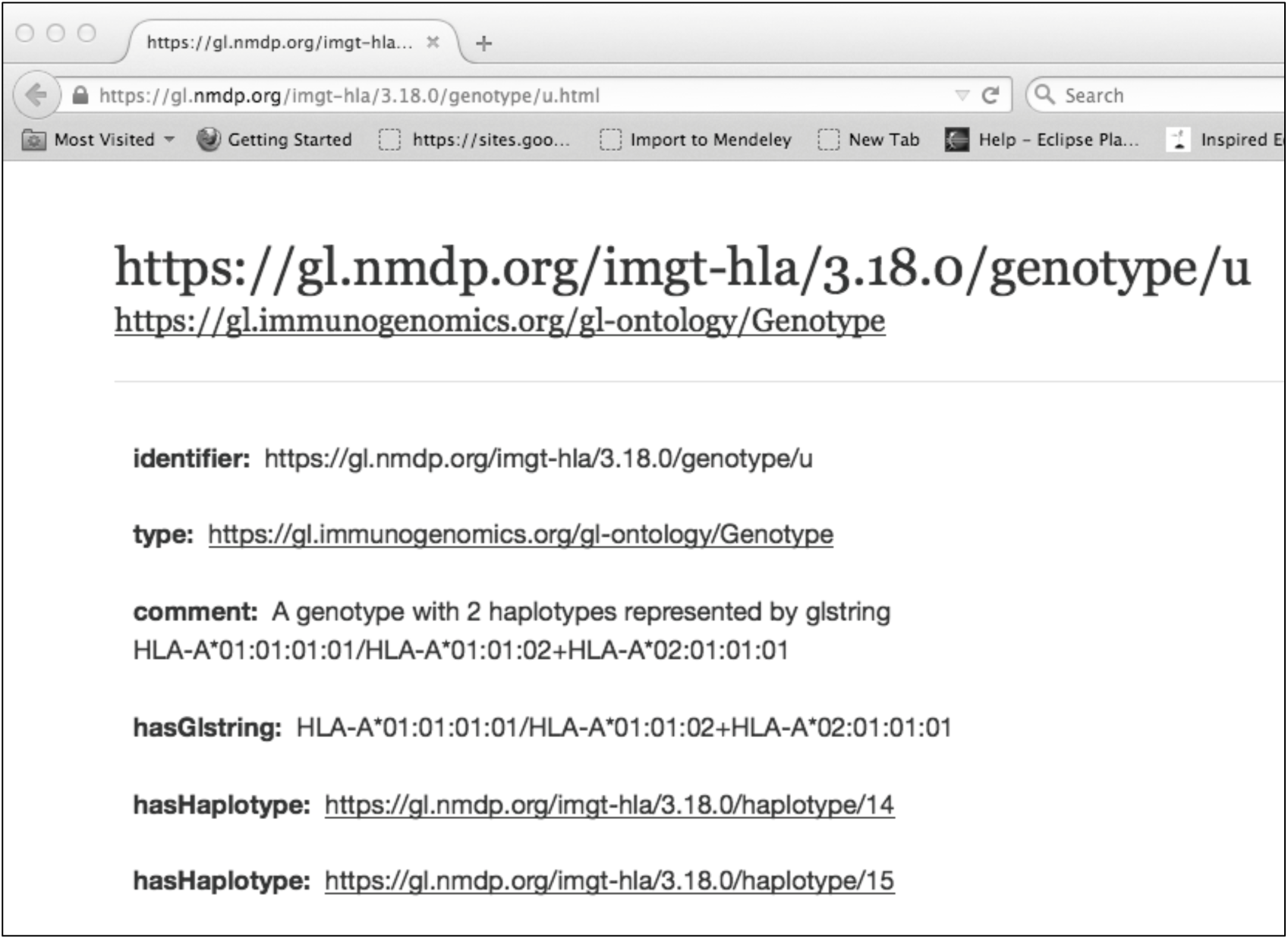
An HTML view of the URI https://gl.nmdp.org/imgt-hla/3.18.0/genotype/u which represents the GL String HLA-A*01:01:01:01/HLA-A*02:01:01:01+HLA-A*02:01:01:01

#### 3.2.2. Client libraries

For integration into larger software systems, a Java client library is provided that wraps HTTP interaction with the RESTful web service API behind simple method calls.

Code listing (excerpt) GlClient.java:

~~~
public interface GlClient {
  String registerAlleleList(String glstring) throws GlClientException;
  AlleleList getAlleleList(String uri) throws GlClientException;
  AlleleList createAlleleList(String glstring) throws GlClientException;
  …
}
~~~

The register methods accept a GL String representation and return the URI of the newly registered GL resource. The get methods accept a URI identifier and return the e.g., AlleleList object representation of the allele list identified by that URI, if such exists. The create methods perform a call to the register method followed by a call to the get method, so for an e.g., allele list in GL String representation the caller receives an AlleleList object representation of the newly registered allele list resource.

Usage:

~~~
GlClient client = …;
~~~

~~~
Allele a0 = client.getAllele(“https://gl.nmdp.org/imgt-hla/3.18.0/allele/0”);
Allele a1 = client.getAllele(“https://gl.nmdp.org/imgt-hla/3.18.0/allele/1”);
AlleleList alleleList0 = client.createAlleleList(a0, a1);
~~~

~~~
String uri = client.registerAlleleList(“HLA-A*01:01:01:01/HLA-A*01:01:01:02N”);
AlleleList alleleList1 = client.getAlleleList(uri);
~~~

Note the client library caches frequently used resources in order to speed up consequent queries.

The client library (and by extension, the command line tools described below) can be configured to use either the application/json media type or the application/xml media type when accessing the Genotype List service RESTful web service APIs.

The JSON representation uses a linked data style format, where associations between parent and child resources are described using URI identifiers. Thus a single request for a high-level GL resource type, such as multilocus unphased genotype, may result in several separate queries against the service to populate the full hierarchy of resources. The response to each individual request will be relatively small in content size.

The XML representation uses a fully specified nested object representation, where associations between parent and child resources are contained as nested XML elements in a single file. This results in only a single request to the service for a high-level resource type. The response to the request will be much larger in content size, however.

This configuration choice allows the user to optimize for response content size or number of requests as necessary.

#### 3.2.3. Command-line tools

For straightforward integration into scripting applications such as typing report submission or sequence analysis pipelines, a suite of command line tools is provided. These command line tools wrap the client library and allow for batch processing of large numbers of GL String resources at one time.

One command line tool for each GL resource type is available. Resources in GL String representation are read via the standard input stream (stdin) or an input file. The URI identifiers are written to the standard output stream (stdout) or an output file.

For example,

~~~
$ gl-register-allele-lists \
  --namespace https://gl.nmdp.org/imgt-hla/3.18.0/ \
  --glstrings glstrings.txt \
  --identifiers identifiers.txt
~~~

reads allele lists in GL String representation from the file glstrings.txt, registers them against the instance of Genotype List Service at the namespace https://gl.nmdp.org/imgt-hla/3.18.0/, and writes the URI identifiers of the newly created allele list resources to the file identifiers.txt.

### 3.3. Accessory services

As a low-level utility service, the GL Service provides a robust building block on which to create accessory services. Two such services are already available, a service that provides the means to lift over a GL resource defined in one version of the IMGT/HLA allele reference database to an equivalent GL resource defined in another version, and a service that maps allelic ambiguity represented by an allele list GL resource to an arbitrary ambiguity encoding.

#### 3.3.1. Liftover service

Given two or more instances of the GL Service configured with different versions the IMGT/HLA Database nomenclature in strict mode, it may be more straightforward to compare GL resources if they are both registered in the same instance of the GL service.

A liftover service has been implemented that given a source namespace, a URI identifying a GL resource in the source namespace, and a target namespace will attempt to lift over the source GL resource to the GL Service at the target namespace.

For example, an allele list resource identified by the URI https://gl.nmdp.org/imgt-hla/3.17.0/allele-list/0 with GL String representation “HLA-A*01:01:01:01/HLA-A*01:01:01:02N” may lift over to the target namespace https://gl.nmdp.org/imgt-hla/3.18.0/ via the liftover service, which if successful will return the URI identifying the newly created allele list in the target namespace, e.g. https://gl.nmdp.org/imgt-hla/3.18.0/allele-list/0.

Like the GL Service, the liftover service provides RESTful web service APIs.

#### 3.3.2. Ambiguity service

Many legacy or arbitrary encodings of allelic ambiguity are in use in the HLA and KIR communities (e.g., NMDP Allele Codes [18]). To adapt these encodings to allele list GL resources registered in instances of the GL Service configured with the IMGT/HLA Database nomenclature in strict mode, an ambiguity service has been implemented. This is a RESTful service for assigning a short name to an allele list representing allelic ambiguity. This service could also be used for serology mappings, epitope group mappings, search determinant mappings, protein level allele name to strict-mode IMGT/HLA nomenclature mappings, P group mappings, etc.

## 4. Discussion

While standards are critical for HLA data interoperability, they are not meaningful until useful tools are developed and made available for community use. Here we have described the development of web services to create and retrieve complete HLA typing data in standardized formats.

The tools being developed here provide the HLA researcher, clinician and lab technician a common resource for managing HLA data in a standardized fashion. We envision these tools augmenting workflows through creation of new instances of HLA typing objects when needed, and retrieval of those objects and their associated metadata when called upon. By exchanging URIs and dereferencing them through the GL Service, users can easily transmit HLA genotypes in a variety of useful formats (GL String, HTML, XML, JSON, OWL/RDF, N3 and QR Code). The GL Service may be combined with other tools and services to create useful workflows for different use cases, such as sending genotyping data to registries, or research pipelines. They may be used as persistent identifiers for genotyping results in publications, and exchanged between research collaborators, and integrated with semantic web genotype-phenotype and disease association resources.

While full allele and genotype ambiguity can be recorded into GL Strings, these strings can be quite large and legacy systems with limited data formats in their transport mechanisms may not be able to accommodate them. Newer formats for HLA typing (e.g., HML 1.0) and clinical data exchange (e.g., HL7 CDA, HL7 FHIR) often provide mechanisms to include pointers to data rather than the data itself. The GL Service is well suited for this model of clinical reporting. Here, the GL String would be registered in the GL Service and the URI may be used in a genotyping report, perhaps with a QR code included in a printed document if needed. Using separate service to register the methodology of the lab test (e.g., NCBI Genetic Testing Registry), the IDs may be sent to the registry, which can then dereference them in their systems. This example is illustrated in Figure 8.

**Figure 8.**
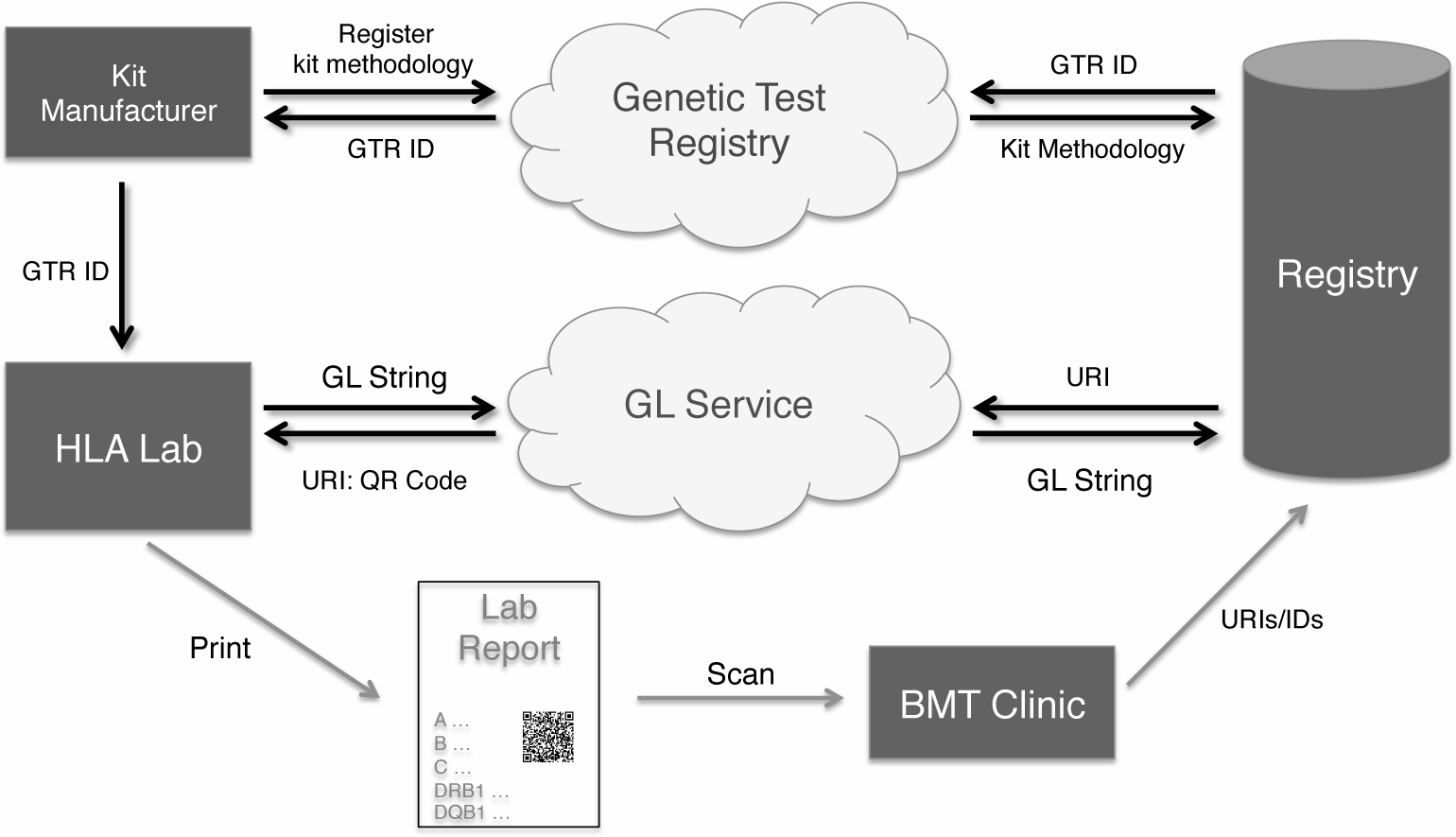
Using the GL Service with a donor registry.

The QR Code format could be used to augment a current HLA laboratory typing report so make it machine readable. By encoding the data using the GL Service and obtaining a QR Code included with the report the recipient of the typing report might then use their smart phone or desktop computer to decode the URI identifier in the QR Code and then retrieve the full ambiguity in GL String representation using the Genotype List service.

Toward these ends, tools leveraging the GL Service to store and exchange genotype data are under development. For example, the Toolkit for Immunogenomic Data-Exchange and Storage (TIDES), accepts genotype data generated by a variety of platforms (e.g., HLA Fusion and Conexio Genomics Assign ATF), generates compact, standardized GL Strings for these data, registers them with the GL Service, disambiguates strings, provides reports in a variety of formats, and traffics formatted genotype data directly to data-analysis software. The use of TIDES and other applications that leverage the GL Service will foster the sharing, reuse, revaluation and meta-analysis of genotype data in ways that have not previously been possible.

## 5. Acknowledgements

This work was supported by Office of Naval Research (ONR) grant N00014-12-1-0142 (MM and RPM), National Institutes of Health (NIH) grants U01AI067068 (SJM and JAH), awarded by the National Institute of Allergy and Infectious Disease (NIAID), and R01GM109030 (SJM, JAH, MM, and RPM), awarded by the National Institute of General Medical Sciences (NIGMS). The content presented is solely the responsibility of the authors and does not necessarily represent the official views of the NIH, NIAID, NIGMS, ONR, Department of Office of Naval Research, the Department of the Navy, the Department of Defense, or the US Government.

## References

[1] Hollenbach JA, Mack SJ, Gourraud P-A, Single RM, Maiers M, Middleton D, et al. A community standard for immunogenomic data reporting and analysis: proposal for a STrengthening the REporting of Immunogenomic Studies statement. Tissue Antigens 2011;78:333–44.

[2] Milius RP, Mack SJ, Hollenbach J a, Pollack J, Heuer ML, Gragert L, et al. Genotype List String: a grammar for describing HLA and KIR genotyping results in a text string. Tissue Antigens 2013;82:106–12.

[3] Berners-Lee T, Fielding RT, Masinter L. Uniform Resource Identifier (URI): Generic Syntax. IETF RFP 3986 (standards Track), Internet Eng Task Force 2005.

[4] Fielding RT. Architectural styles and the design of network-based software architectures. University of California, Irvine, 2000.

[5] Fielding RT, Taylor RN. Principled design of the modern Web architecture. ACM Trans Internet Technol 2002;2:115–50.

[6] Robinson J, Halliwell JA, McWilliam H, Lopez R, Parham P, Marsh SGE. The IMGT/HLA Database. Nucleic Acids Res 2013;41:D1222–7.

[7] W3C OWL Working Group. OWL 2 Web Ontology Language: Document Overview (Second Edition)- W3C Recommendation 11 December 2012 2012.

[8] Gandon F, Schreiber G. RDF 1.1 XML Syntax - W3C Recommendation 25 February 2014, http://www.w3.org/TR/2012/REC-owl2-overview-20121211/, last accessed 18 March 2015

[9] Peterson D, Gao S, Malhotra A, Sperberg-McQueen CM, Thompson HS. W3C XML Schema Definition Language (XSD) 1.1 Part 2: Datatypes - W3C Recommendation 5 April 2012, http://www.w3.org/TR/xmlschema11-2/, last accessed 18 March 2015

[10] Gao S, Sperberg-McQueen CM, Thompson HS. W3C XML Schema Definition Language (XSD) 1.1 Part 1: Structures - W3C Recommendation 5 April 2012, http://www.w3.org/TR/xmlschema11-1/, last accessed 18 March 2015

[11] DeRose S, Maier E, Orchard D, Walsh N. XML Linking Language (XLink) Version 1.1 - W3C Recommendation 06 May 2010, http://www.w3.org/TR/xlink11/, last accessed 18 March 2015

[12] MySQL, http://www.mysql.com, last accessed 18 March 2015

[13] Oracle, http://www.oracle.com, last accessed 18 March 2015

[14] Protégé - A free open-source ontology editor and framework for building intellegent systems, http://protege.stanford.edu/, last accessed 18 March 2015

[15] Apache Jena - A free and open source Java framework for building Semanic Web and Linked Data applications, //jena.apache.org/, last accessed 18 March 2015

[16] RESTClient, a debugger for RESTful web services, https://addons.mozilla.org/en-us/firefox/addon/restclient/, last accessed 18 March 2015

[17] RESTEasy, http://resteasy.jboss.org/, last accessed 18 March 2105

[18] Maiers M, Hurley CK, Perlee L, Fernandez-Vina M, Baisch J, Cook D, et al. Maintaining updated DNA-based HLA assignments in the National Marrow Donor Program Bone Marrow Registry. Rev Immunogenet 2000;2:449–60.

